# Ascorbic acid derived carbon dots promote circadian rhythm and contribute to attention deficit hyperactivity disorder

**DOI:** 10.1101/2022.03.01.482578

**Authors:** Jian Huang, Yun Wang, Zhaomin Zhong, Yurong Ma, Changhong Liu, Keru Deng, Hui Huang, Yang Liu, Xin Ding, Zhenhui Kang

## Abstract

Attention deficit hyperactivity disorder (ADHD) is one of the most prevalent psychiatric disorders in children, and ADHD patients always display circadian abnormalities. While, the ADHD drugs currently used in clinic have strong side effects, such as psychosis, allergic reactions and heart problems. Here, we demonstrated carbon dots derived from the ascorbic acid (VCDs) could strongly rescue the hyperactive and impulsive behaviour of a zebrafish ADHD disease model caused by *per1b* mutation. VCDs prolonged the circadian period of zebrafish for more than half an hour. In addition, the amplitude and circadian phase were also changed. The dopamine level was specifically increased, which may be caused by stimulation of the dopaminergic neuron development in the midbrain. Notably, it was found that the serotonin level was not altered by VCDs treatments. Also, the gene transcriptome effects of VCDs were discussed in present work. Our results provided the dynamic interactions of carbon dots with circadian system and dopamine signaling pathway, which illustrates a potential application of degradable and bio-safe VCDs for the treatment of the attention deficient and hyperactive disorder through circadian intervention.

**Brief summary:** 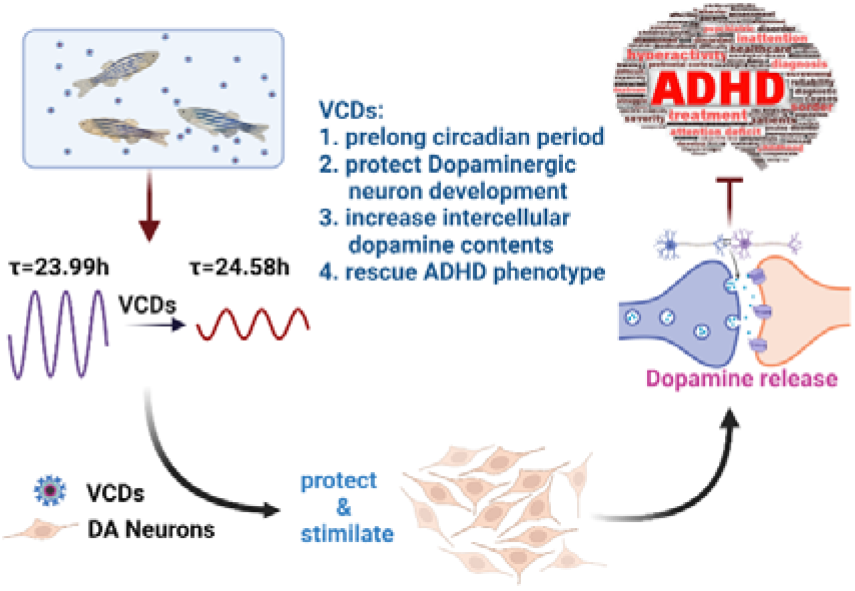

## 1. INTRODUCTION

Attention–Deficit/hyperactivity Disorder (ADHD) is one of the most prevalent and highly heritable psychiatric disorders in childhood^1 2^, affecting about 5-7% of the population^3^, and the number has been continuously increasing these years. ADHD is characterized by hyperactivity, impulsivity and inattention^4^, which impairs normal development of social and academic functions in children, and usually leads to secondary problems, such as drug addiction in adults^5^. The most commonly used drug in the treatment of ADHD is methylphenidate, also called Ritalin, which acts as an indirect agonist on dopaminergic synapses^6^. However, it has many side effects including psychosis, allergic reactions, and heart problems^7^. It also has substance abuse effect similar to amphetamine (ice narcotics) and cocaine^8^.

Zebrafish (Danio rerio) has been validated as a good model for studying the developmental bases of behaviors and drug screening as well^9^. We previously showed that zebrafish key circadian gene per1b mutant shows typical ADHD phenotypes such as hyperactivity, impulsivity and learning deficit, and can be fully rescued by commercial ADHD drug, Ritalin and another monoamine oxidase inhibitor, deprenyl. Significantly, the dopamine level was also decreased in per1b mutant^10^ and it was widely accepted that the *per1b* mutant serves as a perfect ADHD animal model^11^.

In 2016, Xue et.al. reported that aggregated single-walled carbon nanotubes can inhibit methamphetamine self-administration by facilitating dopamine oxidation^12^. Donghoon et.al reported a graphene quantum dots which can prevent α-synucleinopathy in Parkinson’s disease through protecting against dopamine neuron loss^13^. All these findings strongly enlighten us that carbon-based nanomaterial may have the potential application in drugs study. However, some essential properties still need to be considered, for instance, biosafety and bioeffect-reversibility.

Carbon dots (CDs) have become a rising star in the Nano-carbon family due to their excellent and relative biosafety characters^14^, which were widely used in Nano electronics^15^, biosensors^16^ and drug delivery systems^17^. Recent studies showed that a surface modified CDs can specifically deliver dopamine hydrochloride into midbrain and target to neuron diseases^18^. We have previously reported a degradable CDs which can be totally degraded into H_2_O and CO_2_^19^.

Given the important roles of dopamine in the regulation of locomotion and motivational behaviors such as ADHD, we hypothesized that CDs might be useful for the treatment of ADHD by a similar mechanism related to the dopamine system. Here, we show that the degradable and bio-safe CDs synthesized by ascorbic acid (VCDs) specifically stimulate dopamine secretion and dopaminergic neuron development in zebrafish, and can strongly reverse the hyperactivity and impulsivity phenotypes of the ADHD animal model, *per1b* mutant. These results give a new version of CDs application and provide a clue to support the development of new therapeutic agents for abnormal psychiatric and neurological disorders.

## 2. EXPERIMENTAL SECTION

### 2.1 Materials

All procedures were approved by the Soochow university animal care and use committee and were in accordance with governmental regulations of China.

### 2.2 VCDs fabrication

The fabrication of VCDs were developed using the bottom-up method as described by Li et.al.,20 with a little modification which the ultra-water described in Li’s paper was replaced by ascorbic acid solution8.

### 2.3 VCDs characterization

The transmission electron microscopy (TEM) and high-resolution TEM (HRTEM) images were obtained with a FEI/Philips Tecnai G2F20 TWIN TEM. The Fourier transform infrared (FT-IR) spectra of the carbon dots were obtained with a Bruker Fourier Transform Infrared Spectrometer (Hyperion). The photoluminescence (PL) study was carried out on a Horiba Jobin Yvon (Fluoro Max-4) Luminescence Spectrometer, whereas the UV-vis spectra were obtained by a PerkinElmer UV-vis spectrophotometer (Lambda 750).

### 2.4 Cellular toxicity test

CCK8 assay was used to detect the cytotoxicity of VCDs to animal cells. MN9D, the mice midbrain dopaminergic neuron cell line, was treated with increasing concentration series (2.5, 5, 10 and 20 μg/mL) of VCDs for 24 hours. Then we analyzed cell survival by CCK8 assay. All of the experiments were performed for three statistical repeats and three biological repeats. For toxicity test on zebrafish larvae, different concentrations of VCDs were directly added into system water containing 50 3dpf larvae for at most 7days. We calculated the survival rate of larvae every day.

### 2.5 Fish husbandry and embryo production

Zebrafish AB strain was raised according to the standard protocols. Embryos were produced by pair mating, and were raised at 28.5 °C in E3 embryo medium. For RNA isolation, embryos of different treatments were collected and stored at −80 °C. For immunofluorescence staining, larvae were fixed in 4% paraformaldehyde (PFA) in PBS for 3 h at room temperature (RT) or overnight at 4 °C, then washed briefly with PBS, dehydrated, and stored in 100% methanol at −20 °C until use.

### 2.6 Locomotor activity assay

Locomotor activity analysis was carried out as described previously10. Different genotypes of male adult fish were placed in 1 L tanks (20*8.5*6 cm) and transferred to the experiment room. After 1 h of acclimation, the VCDs were added in the system water with final concentration of 2.5 μg/mL for ten minutes, and swimming activities were measured in an automated video-tracking system (Videotrack, Viewpoint life sciences), the movement of each fish was recorded and analyzed using Zebralab4.10 software. The track of fish swimming was also recorded automatically.

### 2.7 Mirror-image attack test

The same tank in the locomotor activity assay for adult fish was used, except that an 8*8 cm mirror was added outside one side of the tank. The fish was treated with 2.5 μg/mL VCDs for ten or 30 minutes, and then transferred into the test tank immediately. The numbers of time adult fish attacked the side with mirror were recorded and analyzed.

### 2.8 Measurement of neurotransmitters and amino acid contents

Embryos were treated with 2.5 μg/mL VCDs or vitamin C for 24 hours from 96hpf (hours post fertilization) to 120hpf, using system water as control. 100 mg of larvae (about 100 individuals) were fast frozen and were stored at −80 °C until use. The neurotransmitters contents were measured with UPLC-QQQ-MS (Waters Acquity UPLC, AB SCIEX 5500 QQQ-MS). All standard samples were purchased form Sigma Aldrich Company.

### 2.9 Quantitative real-time PCR

Total RNAs were extracted from 50 5dpf larvae treated with different concentrations of VCDs by Trizol reagent (ambion). After treated with DNase, 3 μg of purified total RNAs were reversely transcribed into cDNAs. Quantitative real-time PCR (qRT-PCR) was performed in an ABI StepOnePlus instrument with the SYBR green detection system (Invitrogen). PCR thermal profiles were 40 cycles with 10 s at 95 °C and 30 s at 60 °C each cycle. Experiments were performed in triplicate, each with three different biological samples (nine replicates). All results were normalized to the expression level of the housekeeping gene β-actin. qRT-PCR results were shown as a relative expression level calculated by the 2-ΔΔCT method21. p values were calculated with one-way ANOVA test or Student’s t test.

### 2.10. Immunofluorescence staining with tyrosine hydroxylase antibody

5-dpf zebrafish larvae were used to perform immunofluorescence staining. Fixed larvae were merged in 30% sucrose in PBS overnight and then transferred to Tissue-Tek O.C.T. and stored at −26 °C. Sections with a thickness of 14 μm were obtained with a cryostat (Leica CM1850). Immunofluorescence was performed as follows: slides with sections were washed with PBS (3X, each for 10 min), followed by PBST (3X, each for 10 min). Sections then were blocked in 3% bovine serum albumin in PBS for at least 30 min and incubated with TH primary antibody (mouse, 1:1000; Immunostar) overnight at 4 °C. The following day, sections were washed with PBS (6X, each for 10 min). And then incubated with secondary antibody (anti-mouse Alexa 488; A21202, Invitrogen) for 2 h, followed by washing with PBS (6X, each for 10 min). Slides were cover slipped using DAPI solution (Vector Lab, H1200). Images were taken using a Zeiss compound microscope (Axio Imager M2) with a digital camera and processed with Adobe Photoshop CS.

### 2.11 Transcriptome analysis of VCDs treated fish

96hpf larvae were treated with 2.5 μg/mL(LC), 5 μg/mL(MC) and 10 μg/mL(HC) of VCDs respectively for 24 hours. Around 50 larvae were harvested and stored at −80 □ until use. Total RNA was extracted using RNeasy Mini kit (Qiagen, 74104), the purity and concentrations of extracted RNA were estimated with a Nanodrop 2000 spectrophotometer (Thermo Fisher). Total amount of 500 ng per sample was used for the RNA sample preparations. Sequencing libraries were generated using NEBNest-Ultra RNA library prep kit (NEB, USA) following the manufacturer’s instructions. The libraries were sequenced on an Illumina Novaseq 6000 platform. Transcriptome assembly was performed according to the former described elsewhere^22^.

## 3. RESULTS AND DISCUSSION

### 3.1 Characterization of VCDs

The VCDs were synthesized by one-step electrochemical treatment of ascorbic acid aqueous solution. The morphologies of VCDs were obtained by transmission electron microscopy (TEM). As shown in Figure 1A, TEM images revealed that VCDs were well dispersed, and their diameter ranged from 2 to 10 nm, with average size about 5 nm. These VCDs can photodegrade themselves under visible light ^19^. Figure 1B showed an obvious decrease in diameter distributions after 120 h of degradation process ^19^. The UV-vis absorption spectra (Figure S1A) and Fourier transform infrared spectra (Figure S1B) of VCDs further suggested that VCDs possess aromatic sp^2^ domains, multi-conjugate C=O/C-O structures and rich oxygen functional groups ^19^. The high-resolution XPS spectrum of C 1s (Figure 1C) exhibited three main peaks, in which the anterior peak located at 284.8 eV was attributed to the bonding structure of sp^2^ C, whereas the other two peaks located at 286.3 and 288 eV corresponding to the C-O and C=O bonds, respectively ^19^.

**Figure 1.**
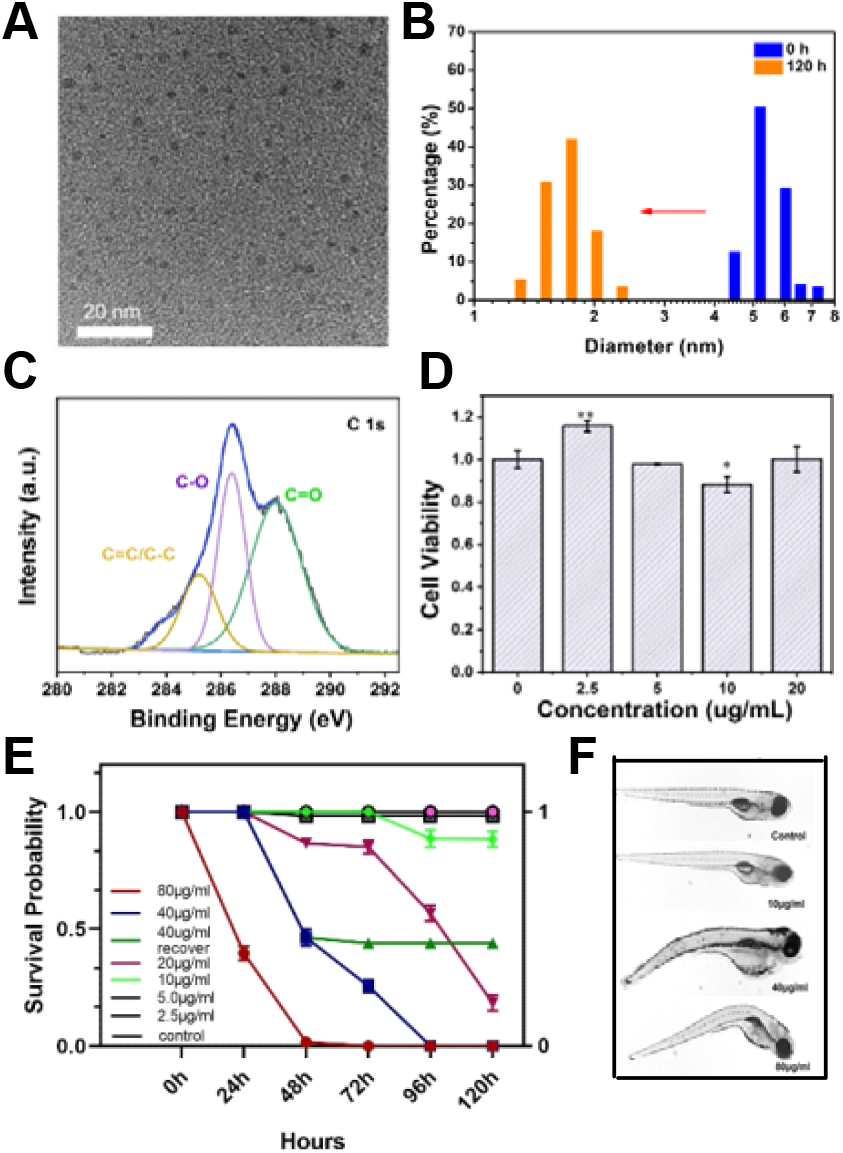
Structure and properties of VCDs. A, TEM images of VCDs. B, diameter distribution of VCDs. C, High-resolution C 1s XPS spectra of VCDs. D, cell viability analysis of dopaminergic neuron cell line MN9D treated by different concentrations of VCDs for 24 h, estimated by Cell Counting Kit-8 (CCK8) assay. E, survival rate of VCDs treated zebrafish larvae. F, phenotypes of some of the misshapen larvae.

### 3.2 Cytotoxicity analysis of VCDs

To test the cytotoxicity of the VCDs, we performed CCK8 assay in a mice midbrain dopaminergic neuron cell line MN9D. Cells were treated with different concentrations of VCDs for 24 hours. As shown in Figure 1D, 2.5 μg/mL VCDs could significantly increase the proliferation of MN9D cells for about 10% compared with the controls. In addition, the cell viability was almost 100% even treated with 20 μg/mL VCDs, indicating low cytotoxicity and excellent biocompatibility. We further tested the toxic effects of VCDs on zebrafish larvae. 3dpf larvae were treated with different concentrations of VCDs for at most 4 days and the survival rate was calculated. As shown in Figure 1E, there were no effects on the survival of zebrafish larvae at concentrations of 2.5 μg/mL and 5 μg/mL, and only one misshapen larvae was detected within totally 58 larvae treated with 10 μg/mL VCDs, as shown in Figure 1E and F. However, when the concentration came to 80 μg/mL, almost all the larvae (52/60) showed sever developmental malformation and were died within 48 hours (Figure1E and F). To test whether this kind of effect can be reversed or not, 120 larvae were treated with 40 μg/mL VCDs, and half of them died after 48 hours. Then we divided the survival fish into two groups. One was raised in normal system water for another 72 hours, and the other group was continuously treated with 40 μg/mL VCDs. As shown in Figure 1E, the larvae raised in system water stopped dying and lived well compared with the continuously treated group which were totally died within another 48 hours, indicating that the effects of VCDs can be reversed.

### 3.3 VCDs affects circadian rhythms of zebrafish larvae

By now, we showed VCDs may serve as biosafe materials with bioeffect-reversibility, and then, in the next study, we further investigated its biofunctions on a zebrafish animal model. We continually monitored the locomotor activities from 5dpf to 10dpf under dim light condition (DD) which can reflect the internal circadian periods of zebrafish larvae. As a result, we found that larvae treated with 2.5 μg/mL VCDs displayed an ~0.6-h prolonged circadian period compared with the control larvae (Figure 2A). The locomotor amplitude of the VCDs treated larvae was significantly lower than that of untreated controls under DD conditions (Figure 2B). Furthermore, larvae treated with VCDs showed a phase-delayed phenotype with the average time of one hour under DD conditions (Figure 2C).

**Figure 2.**
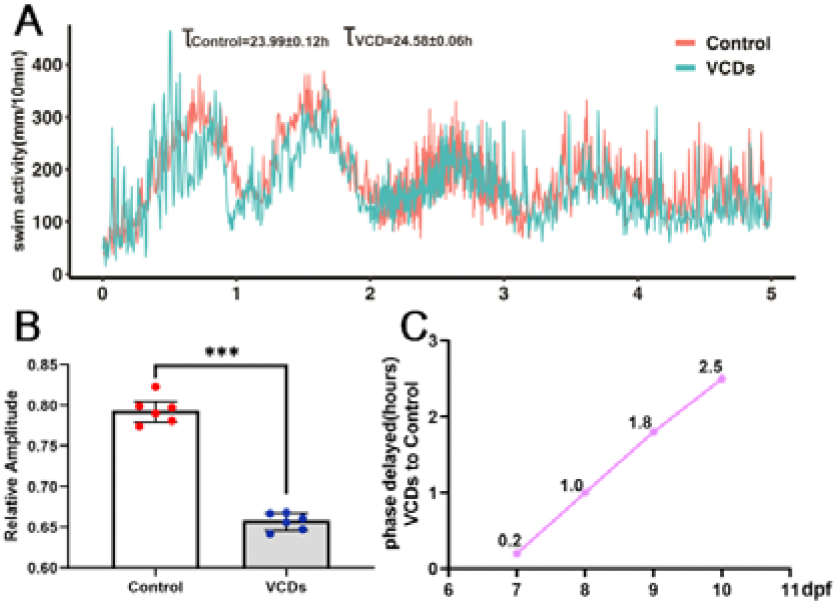
Effects of VCDs on circadian rhythm of zebrafish larvae. A, the locomotor assay of larvae treated by VCDs or not from 5-9 dpf under dim light condition, exhibiting a 0.6 h longer period compared with the untreated controls. Control, n=48, larvae treated by VCDs, n=48. B, the relative amplitude of larvae treated by VCDs was significantly lower than controls DD conditions. control, n= 48; treated larvae, n=48. Student’s t test. C, VCDs displays a 0.8 h phase-delayed effect on zebrafish larvae under DD conditions.

### 3.4 VCDs rescue the ADHD like behaviors

It was reported that aggregated single walled carbon nanotubes can interact with dopamine-related protein in vitro^23^, which may give us a sign that carbon nanomaterial may have a potential influence on the dopamine system. In order to verify this hypothesis, we use zebrafish *per1b* mutant, which was considered as a useful ADHD disease model to perform a series of behavior assay^10–11^. As shown in Figure 3B, the *per1b* mutant was three times more active than the wildtype. When treated with 2.5 μg/ml of VCDs the swimming activity of wildtype fish was not significantly altered. While the *per1b* mutant showed severe decreased activity compared with the untreated *per1b* mutant. In addition, we also found out that the VCDs treatment can recover the less centrophobic phenotype of the ADHD model. In other words, the dramatically increased exploratory behavior which was considered as a reduced anxiety phenotype can be well rescued by the treatment of VCDs (Figure S2, rectangle regions). We also tested the swimming speed of adult fish in different time series under VCDs treatments. As illustrated in Figure 3C, the *per1b* mutant showed more than two times decreased speed after 10 minutes’ treatment of 2.5 μg/mL VCDs in all the tested time points, while the wildtype showed less sensitive to the VCDs at ZT0 (9:00 am) and ZT12(9:00 pm), despite that the swimming speed was significantly decreased at ZT4(1:00 pm) and ZT8(5:00 pm), indicating that VCDs to some extent may play specific roles in circadian mediated neuro-disorder.

**Figure 3.**
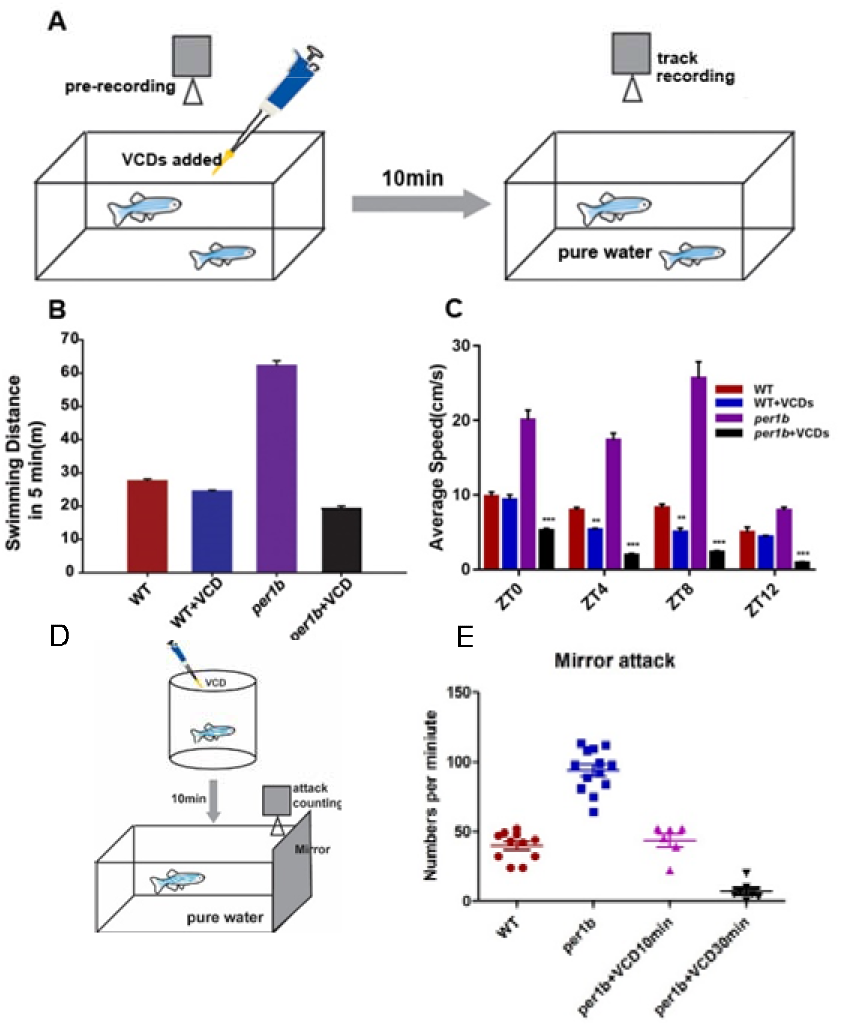
Effects of VCDs on the hyperactivity and impulsivity phenotypes of ADHD zebrafish model (*per1b* mutant). A-C, locomotor assay of wild type (WT) and *per1b* mutant (ADHD model) adult zebrafish treated with 2.5 μg/mL VCDs. A, schematic illustration of hyperactivity rescue experiment. B, the accumulative total tracks in five minutes of the wild-type, per1b mutant, and VCDs treated zebrafish were shown. The hyperactivity phenotype was significantly reversed by VCDs treatments. E, mean average swimming velocities of wild-type, per1b mutant and VCDs treated zebrafish at time -of-day points under normal light-dark conditions. D, schematic illustration of impulsivity rescue experiment. E, impulsive-like behavior of the ADHD model can be rescued by VCDs treatments, estimated by the mirror image attack assay.

We also performed an image-attack assay to estimate the function of VCDs on “perseveration” ^24^, which was another key diagnostic criterion for hyperactivity in ADHD^25^. Adult zebrafish would attack their mirror-image intermittently while ADHD fish showed continuous attack to their self-image, failing to break off. We treated the per1b mutant fish with VCDs for 10 or 30 min, and monitored them with a high-speed camera (Figure 3D). We found that VCDs can efficiently decrease the attack times of *per1b* mutant and the attack phenotype was totally rescued after ten minutes’ treatment (Figure 3E). Thus, the 2.5 μg/mL VCDs can rescue the perseveration-like behavior of ADHD model.

### 3.5 VCDs preferential affect the dopamine signaling pathway

It was reported that abnormal DA system has been implicated in the etiology and pathogenesis of ADHD^26^. It was also reported that altered serotonin level can contribute to the development of ADHD^27^. The plasma amino acids, such as tyrosine and histidine, were correlated with ADHD as well^28^. In order to test whether the rescue of hyperactivity and impulsivity of ADHD model was caused by the alternation of DA system or other neurotransmitters, zebrafish 4dpf larvae were treated with 2.5 μg/mL VCDs for 24 h, and total neurotransmitter contents were measured through target UPLC-QQQ-MS assay. System water and vitamin C solution were used as controls. As shown in Figure 4A, the dopamine level was dramatically increased. In addition, the epinephrine and noradrenaline, the main metabolite of dopamine, was also increased compared with controls (Figure4 B, C). However, the serotonin (5-HT) levels were not changed compared with system water or vitamin C solution (Figure 4D). These results strongly suggested that VCDs may have specific and positive functions on dopamine system. We also measured the plasma amino acid contents. As shown in Figure S3, almost all amino acids except histamine were not altered compared with the water or vitamin C controls, indicating that VCDs may also have effects on histamine related inflammation.

**Figure 4.**
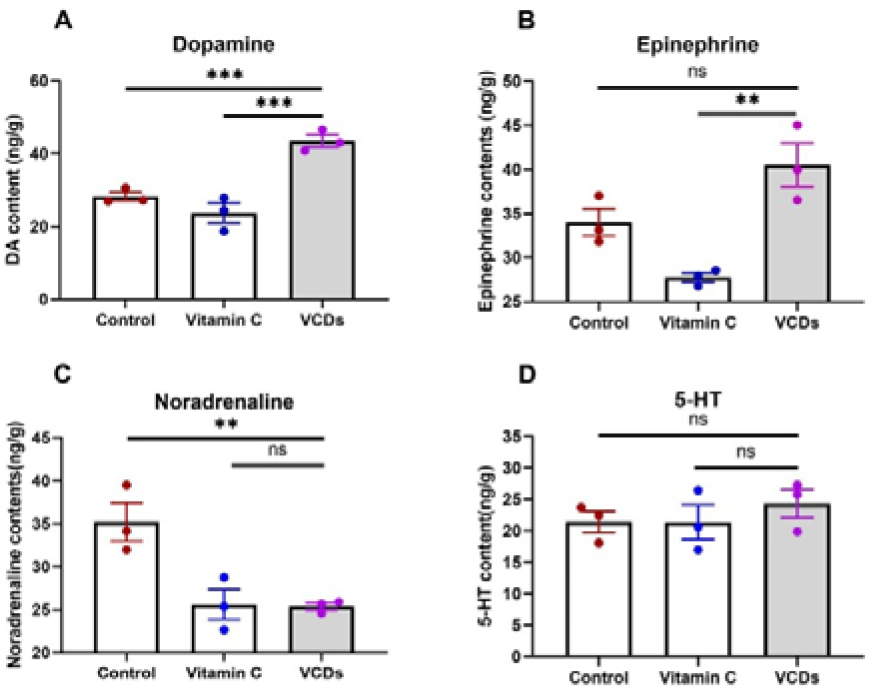
Neurotransmitter’s content of zebrafish 4dpf larvae after treated with 2.5μg/mLVCDs or ascorbic acid for 24 h. A, dopamine. B, Epinephrine. C, Noradrenaline. D, Serotonin (5-HT). Neurotransmitters were determined by UPLC-QQQ-MS. These experiments were performed in triplicate, 50 4dpf larvae were mixed for each measurement. Data were analyzed using one-way ANOVA test, all error bars are ±SEM. ns means not significant.

### 3.6 VCDs affect core circadian and dopamine signaling pathway genes’ expression

We then investigated the effects of VCDs on core circadian genes’ expression levels. As a result, the expression level and patten of *per1a* were not altered (Figure S4A) while *per1b* displayed a 4-hour phase advanced expression patten (Figure S4B). Another two key circadian genes including *clock1a* (Figure S4C) and *bma11b* (Figure S4D) were both upregulated after treated with 2.5μg/mL VCDs. These results indicated that VCDs have important functions on zebrafish circadian regulations.

To investigate whether the VCDs play a role in regulating DA biosynthesis, metabolism, signaling transportation and dopaminergic neuron development, we treated the zebrafish larvae with different VCDs concentrations, and first examined the expressions of dopamine signaling genes. We found that the *dat* (dopamine transporter) was significantly downregulated by the VCDs treatment (Figure 5A), and the *vmat2* (vesicular monoamine transporter) was also downregulated in 2.5 μg/mL and 5 μg/mL VCDs treatments (Figure 5C). However, the *drd2* (dopamine receptor d2) was upregulated in low concentrations of VCDs but downregulated in high concentration of VCDs (Figure 5B), indicating that VCDs may affect the ADHD like behaviors through influencing the dopamine signaling pathways. Since the dopamine level was increased after treated with VCDs, we then tested the expression levels of key dopamine biosynthesis gene, *th* (tyrosine hydroxylase) and found that th gene expression was not changed after treated with 2.5 μg/mL and 5 μg/mL VCDs (Figure 5D). DA could be converted into NE, which was catalyzed by DA β hydroxylase (DBH) and finally converted into epinephrine. We observed that the *dbh* gene expression level was not changed compared with the control (Figure 5E), suggesting that the high contents of dopamine and epinephrine were not caused by the abnormal expression levels of dopamine biosynthesis or related reactions. We also measured the expression level of mao gene, which encodes monoamine oxidases for decomposing DA into DOPAC, and results showed that mao was significantly upregulated (Figure 5F). We speculated that this may be a compensatory response to the high level of dopamine.

**Figure S5.**
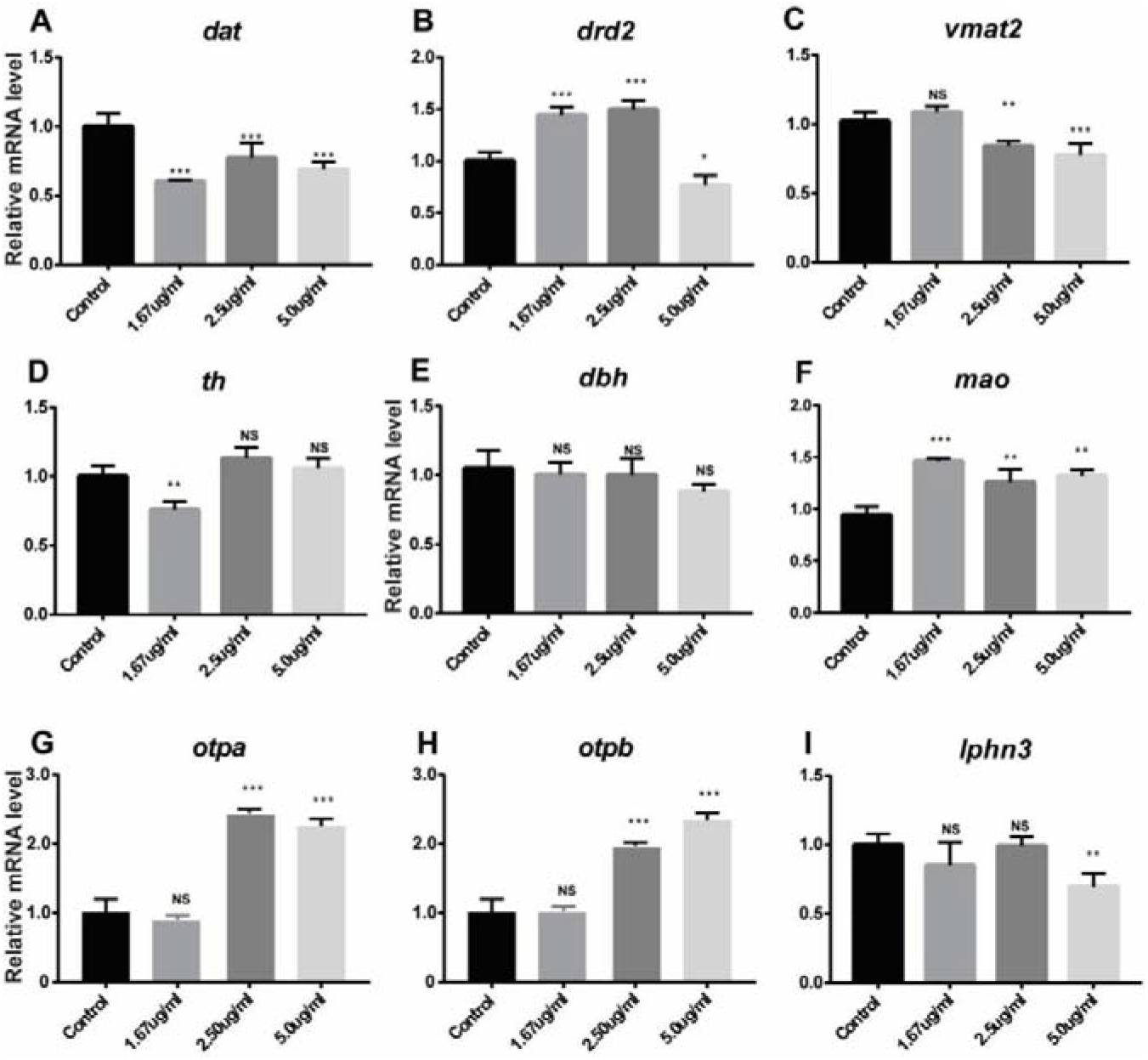
Impaired gene expression levels of dopamine related signaling pathway, shown by QRT-PCR. A-C, dopamine transporter (*dat*; A), dopamine receptor D2 (*drd2*; B) and vesicular monoamine transporter (*vmat*; C). D-F, genes involved in dopamine synthesis and metabolism. Tyrosine hydroxylase (*th*; D), dopamine beta hydroxylase (*dbh*; E) and monoamine oxidase (*mao*; F). G-H, key genes involved in dopaminergic neuron differentiations. Orthopedia homeobox a (*otpa*; G), Orthopedia homeobox b (*otpb*; H). The ADHD susceptible gene *lphn3.1* was not affected by treatment with low and middle concentrations of VCDs. all experiments were performed in triplicate, with three biological samples for each experiment. Data were analyzed using student t test, error bars are ±SEM

### 3.7 VCDs promote dopaminergic neuron development

OTPa (orthopedia homeodomain protein a,) and OTPb (orthopedia homeodomain protein b,) were required for the specification and development of dopaminergic neurons in zebrafish ^29^. Real time PCR showed that both opta and optb were high upregulated after treated with 2.5 μg/mL and 5 μg/mL VCDs (Figure 5 G, H), indicating that VCDs may act via Otp to regulate the specification of dopaminergic neurons. LATROPHILIN3 (LPHN3), encoding a putative adhesion-G-protein-Coupled receptor, was implicated as an ADHD susceptible gene^**26b, 30**^. Here, we found lphn3 was a little bit downregulated only under a relatively high concentration of VCDs (Figure 5I), indicating that VCDs played a tiny role on lphn3.

In zebrafish, dopaminergic neurons in the PT region of ventral diencephalon, projecting into the subpallium, have been implicated in the regulation of complex behaviors^31^, and were concerned as key factors on the pathogenesis of ADHD^26b^. We then performed TH fluorescence immunostaining analysis to see whether the dopaminergic neuron development in the PT region of 4dpf zebrafish larvae was altered or not after treated with VCDs. The dopaminergic neuron in other brain region was also investigated. As shown in Figure 6, compared with the system water control, the VCDs treated brain had more dopaminergic neurons in the PT region. Furthermore, the organization was much more compact under VCDs treatment (Figure 6C, D, F). We also found that the dopaminergic neuron in the eyes was also increased by the VCDs treatments in Figure 6A, B, E. Generally, it had around 5 dopaminergic neurons surrounding the retina. As a contrast, the number was more than 7 in VCDs treated fish. These results suggested that VCDs played a significant role in the whole dopaminergic neuron development. However, the mechanisms still need further investigation.

**Figure 6.**
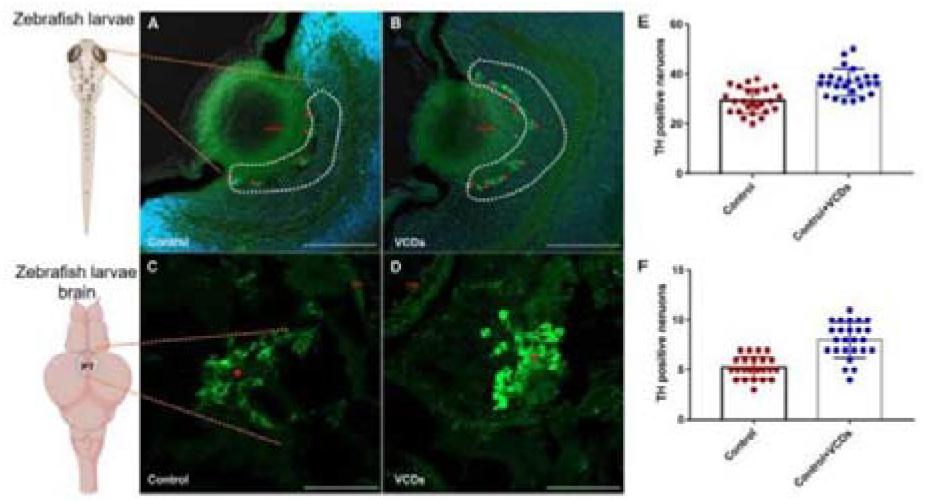
Dopaminergic neurons were significantly increased in zebrafish larvae after treating with VCDs for 24 h. Shown by immunostaining of tyrosine hydroxylase antibody. A, B, E, the dopaminergic neurons were increased in zebrafish eyes, the neurons surrounding the retina was labeled by red arrow. C, D, F, the dopaminergic neurons in the PT of the ventral diencephalon are increased by VCDs treatments. Labeled by the red star. Scale bar, 100 μm. The numbers of dopaminergic neurons were determined with ImageJ in a double-blind manner. N=25 for control larvae and 26 for VCDs treatments. Student’s t test was used in the statistical analysis. Error bars are SEM.

### 3.8 VCDs contribute to cell proliferation tyrosine metabolism and aspartate metabolism

To examine possible effects of VCDs on other processes, we conducted transcriptome sequencing (RNA seq) analysis of zebrafish larvae treated with different VCDs concentrations. We uncovered 177, 160 and 67 downregulated genes in low concentration (1.67 μg/mL), medium concentration (2.5 μg/mL) and high concentration (5 μg/mL) respectively. Among these genes, 12 genes were commonly existed in all three kinds of treatments (Figure 7A). We also discovered 461, 271 and 225 upregulated genes in the three concentrations of treatments as well (Figure 7B), and 83 common genes together with the 12 common downregulated genes were selected to generate an expression heat map, as shown in Figure 7D. Clustering analysis showed that there were more upregulated genes than downregulated genes after VCDs treatment, suggesting that VCDs may act as a transcription activator in zebrafish. Through gene ontology (GO) classification, we found that DEGs (differentially expressed gene) were enriched for the tyrosine metabolism, tyrosine and tryptophan biosynthesis, NOD like receptor signaling pathway, ECM receptor interaction and so on, indicating that VCDs contributed to numerous life processes including stress response, metabolism, and signal transduction in zebrafish. Furthermore, it seemed that VCDs had a preference effect on tyrosine signaling pathways (Figure 7D).

**Figure 7.**
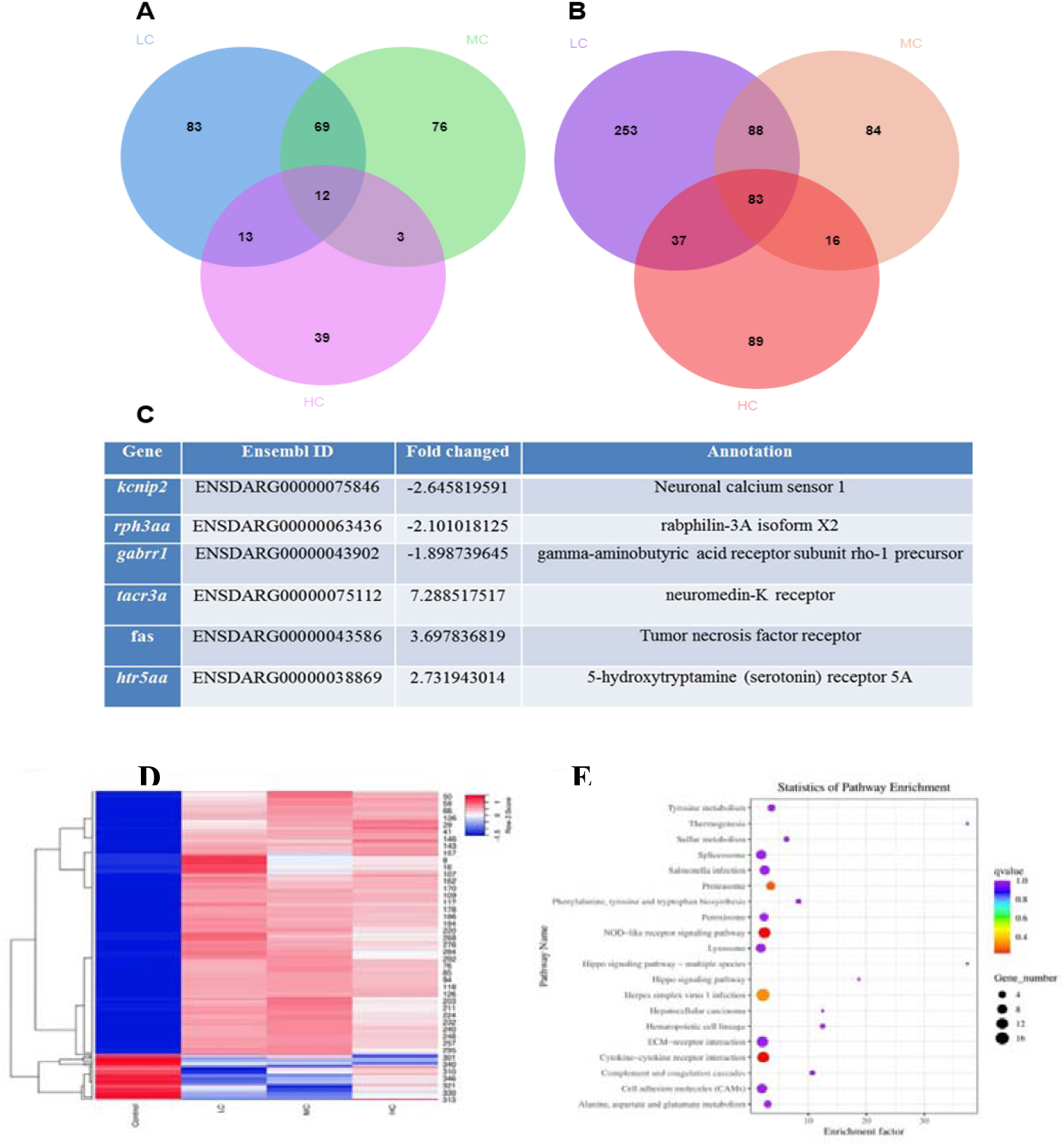
Transcriptome analysis of zebrafish larvae treated with different concentrations of VCDs. A, numbers of downregulated gene in VCDs treated zebrafish. 12 genes were commonly downregulated in zebrafish treated by different concentrations of VCDs. B, numbers of upregulated genes in VCDs treated zebrafish. 83 genes were identified in three different treatments. C, List of genes with significant fold-changes, revealed by transcriptome analysis. The fold-changes were calculated by values of VCDs treated larvae vs. control larvae. Genes with positive values were upregulated after VCDs treatments, and those with negative values downregulated. D, histogram of 95 DEGs in VCDs treated zebrafish. Red colors represent up regulation, and blue colors indicates the down regulation. E, KEGG analysis of the total DEGs. The red three shows the dopamine related gene clusters after VCDs treatments.

## 4. CONCLUSION

This work demonstrated VCDs can strongly rescue the hyperactivity and impulsivity phenotypes of zebrafish ADHD disease model caused by a core circadian gene *per1b* mutation. 2.5 μg/mL treatment of VCDs leading to a 0.6 h prolonged daily period and 0.8 h phase delayed of circadian rhythm. Neurotransmitter content analysis by UPLC-QQQ-MS indicated that the dopamine level in zebrafish larvae was 1.6-fold increased after treated with VCDs, and the epinephrine was also increased by the VCDs treatment, indicating a preference role that VCDs played. Gene expression analysis by real time PCR showed that the dopaminergic development genes including *otpa* and *otpb* were significantly upregulated by the VCDs treatments. Using TH immunostaining, we discovered that the dopaminergic neurons in the PT region of zebrafish brain were highly increased and well organized when treated with VCDs. Moreover, the dopaminergic neurons in the eyes were also increased. Transcriptome sequencing analysis also showed a preference effect on tyrosine (precursor of dopamine) synthesis and metabolism pathways. Our findings may discover a new idea for ADHD therapy, which even can be extended for dopamine related diseases including Alzheimer’s disease and Parkinson disease.

## AUTHOR INFORMATION

### Corresponding Author

Yang Liu* -;

Xin Ding*-

Zhenhui Kang* -.

### Author Contributions

The manuscript was written through contributions of all authors. All authors have given approval to the final version of the manuscript.

## NOTES

The authors declare no competing financial interest.

## ACKNOWLEDGMENT

This work was supported by National Natural Science Foundation of China (31671216, 81871193, 51725204, 21771132, 51972216, 52041202), National MCF Energy R&D Program of China (2018YFE0306105), National Key R&D Program of China (2020YFA0406104, 2020YFA0406101), Innovative Research Group Project of the National Natural Science Foundation of China (51821002), Natural Science Foundation of Jiangsu Province (BK20190041), National college students Innovation and entrepreneurship training program (201710285042Z), Suzhou scientific program (SS202074), Collaborative Innovation Center of Suzhou Nano Science & Technology, the 111 Project, and Suzhou Key Laboratory of Functional Nano & Soft Materials

